# In Silico Modeling of Metabolic State in Single Th17 Cells Reveals Novel Regulators of Inflammation and Autoimmunity

**DOI:** 10.1101/2020.01.23.912717

**Authors:** Allon Wagner, Chao Wang, David DeTomaso, Julian Avila-Pacheco, Sarah Zaghouani, Johannes Fessler, Sequoia Eyzaguirre, Elliot Akama-Garren, Kerry Pierce, Noga Ron-Harel, Vivian Paraskevi Douglas, Marcia Haigis, Raymond A. Sobel, Clary Clish, Aviv Regev, Vijay K. Kuchroo, Nir Yosef

**Author notes:** These authors contributed equally (Allon Wagner, Chao Wang). These authors contributed equally (Vijay K. Kuchroo, Nir Yosef).

## Abstract

Cellular metabolism, a key regulator of immune responses, is difficult to study with current technologies in individual cells Here, we present Compass, an algorithm to characterize the metabolic state of cells based on single-cell RNA-Seq and flux balance analysis. We applied Compass to associate metabolic states with functional variability (pathogenic potential) amongst Th17 cells and recovered a metabolic switch between glycolysis and fatty acid oxidation, akin to known differences between Th17 and Treg cells, as well as novel targets in amino-acid pathways, which we tested through targeted metabolic assays. Compass further predicted a particular glycolytic reaction (phosphoglycerate mutase — PGAM) that promotes an anti-inflammatory Th17 phenotype, contrary to the common understanding of glycolysis as pro-inflammatory. We demonstrate that PGAM inhibition leads non-pathogenic Th17 cells to adopt a pro-inflammatory transcriptome and induce autoimmunity in vivo. Compass is broadly applicable for characterizing metabolic states of cells and relating metabolic heterogeneity to other cellular phenotypes.

## INTRODUCTION

Cellular metabolism is both a mediator and a regulator of cellular functions. Metabolic activities are key in normal cellular processes such as activation, expansion and differentiation, but also play an important role in the pathogenesis of multiple disease conditions including autoimmunity, cancer, cardiovascular disease, neurodegeneration, and aging. Recently, the study of metabolism in immune cells (immunometabolism) has gained particular attention as a major regulator of almost all aspects of immune responses including anti-viral immunity, autoimmunity, and cancer [1]–[9].

Due to the scale and complexity of the metabolic network, a metabolic perturbation may create cascading effects and eventually alter a seemingly distant part of the network, while cross-cutting traditional pathway definitions. Therefore, computational tools are needed to contextualize observations on specific reactions or enzymes into a systems-level understanding of metabolism and its dysregulation in disease. One successful framework has been Flux Balance Analysis (FBA), which translates curated knowledge on the network’s topology and stoichiometry into mathematical objects and uses them to make *in silico* predictions on metabolic fluxes [10]–[13]. FBA methods have proven particularly useful when contextualized with functional genomics data, including gene expression [14].

While such metabolic models aim to represent the behavior of individual cells, their contextualization has generally relied on information collected from bulk population data. However, the advent of single-cell RNA-Seq (scRNA-Seq) has highlighted the substantial extent of cell-to-cell diversity that is often missed by bulk profiles [15], [16], and can be especially prominent in immune cells and associated with their functional diversity [8], [17]–[28]. One of the earliest examples has been the diversity among T helper 17 (Th17) cells [29]. On the one hand, IL-17 producing Th17 cells can be potent inducers of tissue inflammation in autoimmune disorders [30], [31] but on the other hand, these cells are critical in host defense against pathogens [32], [33] and can promote mucosal homeostasis and barrier functions [34]–[36]. Th17 cells with distinct effector functions can be found in patients and animal models and can also be generated in vitro with different combinations of differentiation cytokines, as we have previously demonstrated [34]. We have previously shown that such functional diversity can be captured by studying transcriptional diversity at the single cell level with scRNA-Seq, and enabled the discovery of novel regulators that are otherwise difficult to detect in bulk RNA-Seq analysis [29], [37].

We hypothesized that a similar spectrum of diversity may exist at the immunometabolic level and relate to cell function. However, most cellular assays, including metabolic assays, are normally done in a targeted manner and difficult to undertake at single-cell level. Furthermore, low cell numbers frequently prohibit direct metabolic assays, for example, in the study of immune cells that are present at tissue sites. In contrast, scRNA-Seq, is broadly accessible and rapidly collected across the human body [38], and should allow, in principle, to contextualize metabolic models to the single cell level. A computational method is thus required to systematically address the unique challenges of scRNA-Seq, such as data sparsity, and to capitalize on its opportunities, for example by treating cell populations as natural perturbation systems with a rapidly increasing scale [39].

Here, we present Compass, an FBA algorithm to characterize and interpret the metabolic heterogeneity among cells, which uses available knowledge of the metabolic network in conjunction with RNA expression of metabolic enzymes. Compass uses single cell transcriptomic profiles to characterize cellular metabolic states at single-cell resolution and with network-wide comprehensiveness. It allows detection of metabolic targets across the entire metabolic network, agnostically of pre-defined metabolic pathway boundaries, and including ancillary pathways that are normally less studied, yet could play an important role in the determining cell function [40]. We applied Compass to Th17 cells, uncovering substantial immunometabolic diversity associated with their inflammatory effector functions. In addition to the expected glycolytic shift, we found diversity in amino acid metabolism, and highlighted a unique and surprising role for the glycolytic reaction catalyzed by phosphoglycerate mutase (PGAM) in promoting an anti-inflammatory phenotype in Th17 cells. Compass is a broadly applicable tool for studying metabolic diversity at the single cell level, and its relationship to the functional diversity between cells.

## RESULTS

### Compass — an algorithm for comprehensive characterization of single-cell metabolism

We reasoned that even though the mRNA expression of individual enzymes does not necessarily provide an accurate proxy for their metabolic activity, a global analysis the entire metabolic network (as enabled by RNA-Seq) in the context of a large sample set (as offered by single cell genomics) coupled with strict criteria for hypotheses testing, would provide an effective framework for predicting cellular metabolic status of the cell. This led us to develop the Compass algorithm, which integrates scRNA-Seq profiles with prior knowledge of the metabolic network to infer a metabolic state of the cell **(Figure 1A)**.

**Figure 1.**
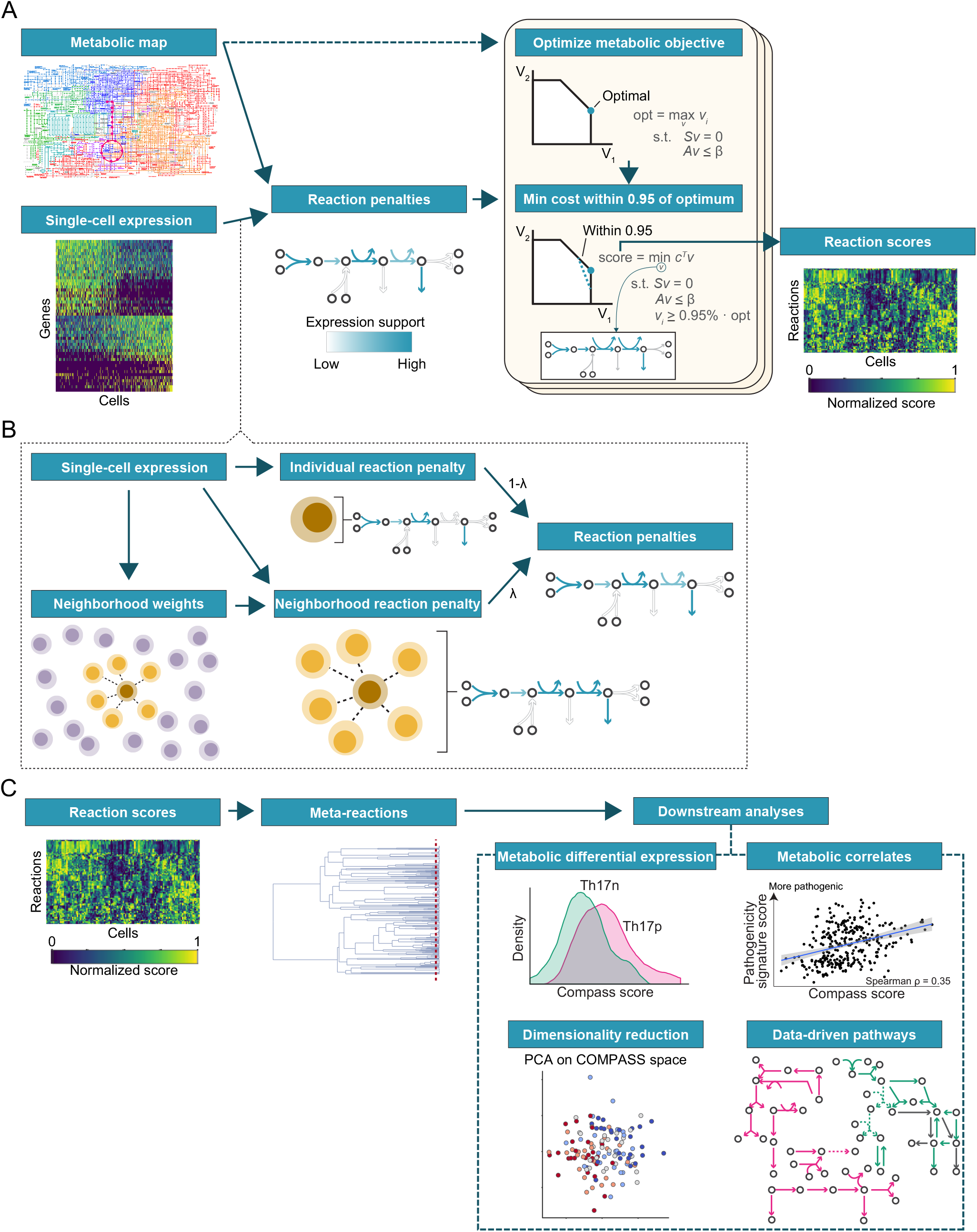
Algorithm overview. (A) Computation of Compass scores matrix. Compass leverages prior knowledge on metabolic topology and stoichiometry (encoded in a GSMM, see main text) to analyze single-cell RNA-Seq expression. Briefly, it computes a reaction-penalties matrix, where the penalty of a given reaction is inversely proportional to the expression its respective enzyme-coding genes. The reactionpenalties matrix is the input to a set of flux-balance linear programs that produce a score for every reaction in every cell, namely the Compass score matrix. (B) To compute the reaction penalties matrix, Compass allows soft information sharing between a cell and its k-nearest neighbors to mitigate technical noise in single-cell library preparation. (C) Downstream analysis of the score matrix. Rows are hierarchically clustered into meta-reactions (agnostically of canonical pathway definitions). The scores are then amenable to common genomics procedures including differential expression of meta-reactions, detecting meta-reactions correlating with a phenotype of interest, dimensionality reduction, and data-driven network analysis (the latter pursued in the accompanying manuscript (Wang et al., biorxiv preprint)).

The metabolic network is encoded in a Genome-Scale Metabolic Model (GSMM) that includes reaction stoichiometry, biochemical constraints such as reaction irreversibility and nutrient availability, and gene-enzyme-reaction associations. Here, we use Recon2, which comprises of 7,440 reactions and 2,626 unique metabolites [41]. To explore the metabolic capabilities of each cell, Compass solves a series of constraint-based optimization problems (formalized as linear programs) that produce a set of numeric scores, one per reaction **(Methods)**. Intuitively, the score of each reaction in each cell reflects how well adjusted is the cell’s overall transcriptome to maintaining high flux through that reaction. Henceforth, we refer to the scores as quantifying the “potential activity” of a metabolic reaction (or “activity” in short when it is clear from the context that Compass predictions are discussed).

Compass belongs to the family of Flux Balance Analysis (FBA) algorithms that model metabolic fluxes, namely the rate by which chemical reactions convert substrates to products and apply constrained optimization methods to find flux distributions that satisfy desired properties (a flux distribution is an assignment of flux value to every reaction in the network) [10]–[13]. In the first step, Compass is agnostic to any measurement of gene expression levels and computes, for every metabolic reaction *r*, the maximal flux 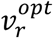 it can carry without imposing any constraints on top of those imposed by stoichiometry and mass balance. Next, Compass relies on the assumption that mRNA expression of an enzyme coding gene should preferably correlate with the flux through the metabolic reaction(s) it catalyzes. It thus assigns every reaction in every cell a penalty inversely proportional to the mRNA expression associated with its enzyme(s) in that cell. Compass then finds a flux distribution which minimizes the overall penalty incurred in any given cell *i* (summing over all reactions), while maintaining a flux of at least 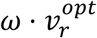 (here *ω* = 0.95) in *r*. The Compass score of reaction *r* in cell *i* is the negative of that minimal penalty (so that lower scores correspond to lower potential metabolic activity).

Using genome-scale metabolic network allows the entire metabolic transcriptome to impact the computed score for any particular reaction, rather than just the mRNA coding for the enzymes that catalyze it. We reasoned that this helps reduce the effect of instances where mRNA expression does not correlate well with metabolic activity, for example due to post-transcriptional or post-translational modifications. This also mitigates the effects of data sparsity, which is characteristic of scRNA-Seq data. The low transcript signal in scRNA-Seq, which results in the extreme case in false-negative gene detections, magnifies the repercussions of sampling bias and transcription stochasticity, and leads to an overestimation of the variance of lowly expressed genes, which in turn leads in turn to false-positive calling of differentially expressed genes [15]. Compass further mitigates data sparsity effects with an information-sharing approach, similar to other scRNA-Seq algorithms [42]–[50]. Instead of treating each cell in isolation, the score vector for each cell is determined by a combined objective that balances the effects in the cell in question with those in its *k*-nearest neighbors (based on similarity of their RNA profiles; here, using *k* = 10; **Figure 1B; Methods**).

The output of Compass is a quantitative profile for the metabolic state of every cell, which is then subject to downstream analyses **(Figure 1C)**. These include finding metabolic reactions that are differentially active between cell types or that correlate with continuous properties of cell state (e.g., expression of a certain group of cytokines). It can also provide unsupervised insights into cellular diversity by projecting cells into a low-dimensional space of metabolic activity (e.g., for visual exploration). The statistical power afforded by the large number of individual cells in a typical scRNA-Seq study adds robustness and allows these downstream analyses to gain biological insight despite the high dimension of the metabolic space in which Compass embeds cells. Finally, because Compass does not rely on a predetermined set of metabolic pathways (or gene sets) such as Reactome [51] or KEGG [52], it allows unsupervised derivation of cell-specific metabolic pathways in a data driven way.

### Th17 cell metabolic diversity reflects a balance between glycolysis and fatty acid oxidation, which is associated with pathogenicity

To demonstrate Compass, we applied it to scRNA-Seq data from Th17 cells, differentiated *in vitro* from naïve CD4+ T into two extreme functional states [34], [53] **(Figure 2A)**. Differentiation with IL-1β+IL-6+IL-23 creates Th17 cells that upon adoptive transfer into recipient mice induce severe neuroinflammation in the form of experimental autoimmune encephalomyelitis (EAE). We refer to these cells as pathogenic Th17 (Th17p). However, when naïve CD4+ T cells are cultured with TGF-β1+IL-6, the resulting Th17 induce only mild-to-none EAE when adoptively transferred to recipient mice. We refer to these cells as non-pathogenic Th17 (Th17n). We performed a Compass analysis of a dataset we generated in a previous study that included 139 Th17p and 151 Th17n cells sorted for IL-17A/GFP+ and profiled using Fluidigm C1 and SMART-Seq2 [29], [37].

**Figure 2.**
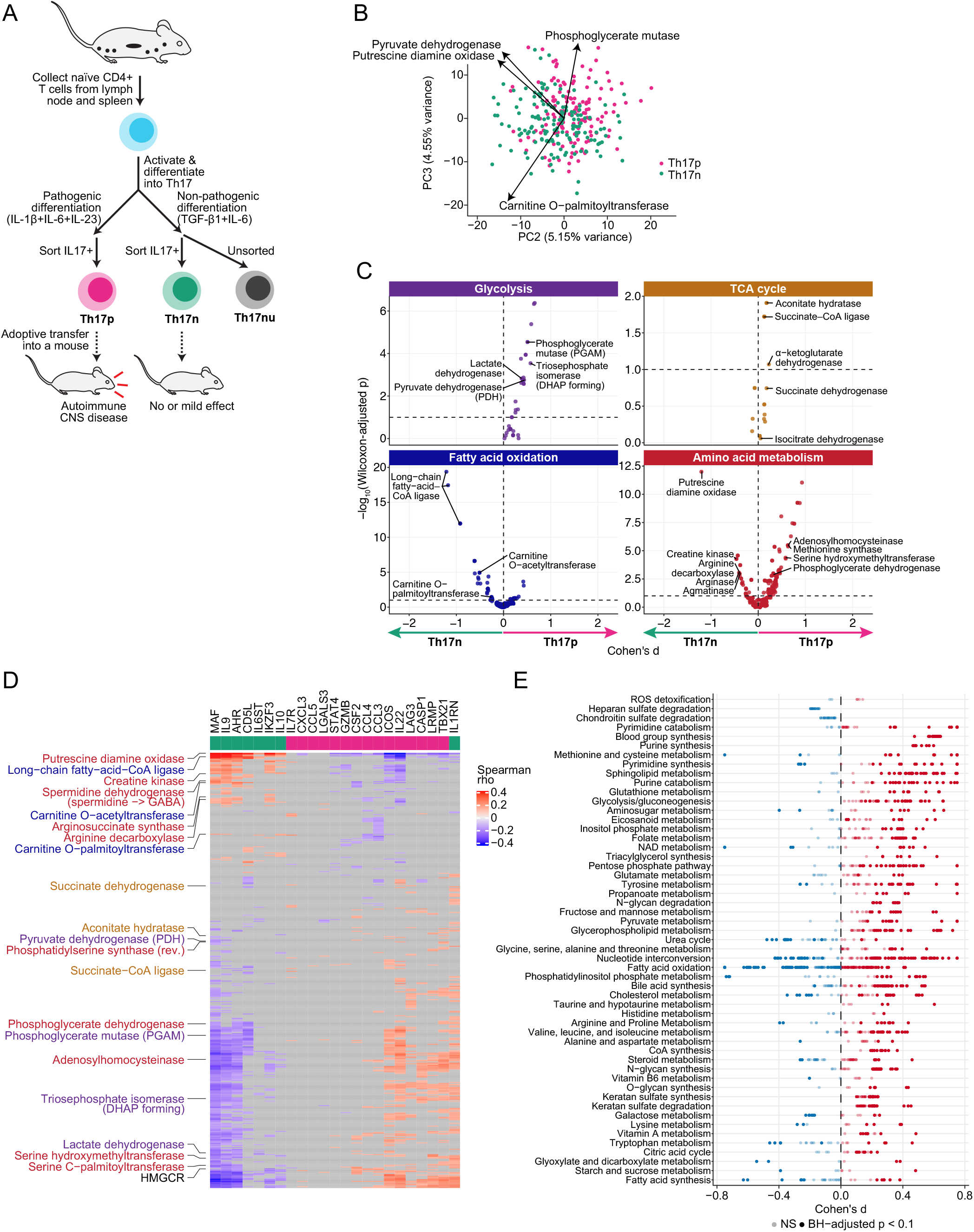
Compass-based exploration of metabolic heterogeneity within the Th17 compartment. (A) The experimental system. Naive CD4+ T cells are collected and differentiated into Th17p or Th17n cells, which are IL-17+ T cells that cause severe or mild-to-none CNS autoimmunity upon adoptive transfer. Th17nu cells are Th17n cells which were not sorted by IL-17 and exhibit higher variability [29]. (B) PCA of the Compass scores matrix (restricted to core metabolism, see main text), with select top loadings shown. (C) Dots represent a single biochemical reactions, Cohen’s d and Wilcoxon rank sum p values computed as described in the main text for a comparison of Th17p vs. Th17n. This computation is done over meta-reactions, and every meta-reaction is expanded into its constituent single reactions (**Methods**), each shown as a separate dot. Only core reactions (**Methods**) are shown. Reactions are partitioned by Recon2 pathways; bottomright panel groups together all Recon2 subsystems associated with amino-acid metabolism. (D) Spearman correlation of Compass scores of single reactions with the expression of pro-pathogenic (magenta) or pro-regulatory Th17n (green) genes [29], none of which is metabolic and therefore none of them directly serves as a Compass input. Only significant correlations (BH-adjusted p < 0.1) are shown in color and non-significant correlation coefficients are greyed out. The rows represent 489 meta-reactions that belong to core pathways (defined as Recon2 subsystems that have at least 3 core reactions), and significantly correlated (or anti-correlated) with at least one of the genes. Key reactions (rows) in pathways discussed in the manuscript are highlighted according to the meta-reaction to which they belong. (E) Dots represent single biochemical reactions, Cohen’s d and Wilcoxon rank sum p values computed as described in panel C. Only core reactions are shown. Reactions are partitioned by Recon2 pathways. Reactions are colored by the sign of their Cohen d’s statistic, and are opaque or transparent according to statistical significance.

We first computed a Compass score for each metabolic reaction in each cell **(Methods)**. We then aggregated reactions that were highly correlated across the entire dataset (Spearman rho ≥ 0.98) into meta-reactions (median of two reactions per meta-reaction; **Supplementary Figure 1**) for downstream analysis.

To investigate the main determinants of Th17 cell-to-cell metabolic heterogeneity, we first analyzed the Compass output as a high dimensional representation of the cells which parallels the one produced by scRNA-Seq, but with features corresponding to metabolic meta-reaction rather than transcripts. We performed principal component analysis (PCA) on the meta-reaction matrix, while restricting it to 784 meta-reactions (out of 1,911) associated with core metabolism (**Methods**) that span conserved and well-studied pathways for generation of ATP and synthesis of key biomolecules.

The first two principal components (PCs) of the core metabolism subspace were associated both with overall metabolic activity and T effector functions (**Figure 2B, Supplementary Figure 2A,B**, **Supplementary Table 1**). PC1 correlated with the cell’s total metabolic activity, defined as the expression ratio of genes coding metabolic enzymes out of the total protein coding genes (Pearson rho = 0.36, p < 4*10^−10^), as well as a transcriptional signature of late stages of Th17 differentiation over time [54] (**Supplementary Figure 2C**, Pearson rho = 0.18, p < 0.003) (**Methods**). PC2 and PC3 represented a choice between ATP generation through aerobic glycolysis versus fatty acid oxidation, which is a prominent finding in immunometabolism while comparing activated Th17 to Tregs, or Teff vs. Tmem cells [3]. Accordingly, they were correlated with multiple Th17 pathogenicity markers, as well as a signature of Th17 pathogenicity consisting of cytokines, chemokines and transcription factors that are associated with each phenotypic group [29], [34] (**Supplementary Figure 2D,E**). PC2 and PC3 were also noticeably associated with nitrogen metabolism, and were enriched in urea cycle targets whose power to modulate Th17 pathogenicity is demonstrated below and in an accompanying manuscript (Wang et al., biorxiv preprint).

### Compass predicts metabolic regulators of Th17 cell pathogenicity

To directly search for metabolic targets that are associated with the pathogenic capacity of individual Th17 cells, we searched for biochemical reactions with differential predicted activity between the Th17p and Th17n conditions according to Wilcoxon’s rank sum p value and Cohen’s d effect size statistics) and defined pro-pathogenic and pro-regulatory reactions as ones that were significantly different in the Th17p or Th17n direction, respectively (**Figure 2C; Supplementary Figure 2F; Methods; Supplementary Table 2**). We next discuss several key predictions, which we follow up on in the rest of the manuscript as well as in the accompanying manuscript (Wang et al., biorxiv preprint).

Metabolic reactions in both primary and ancillary pathways were associated with Th17 cell pathogenicity (1,213 or 3,362 reactions out of 6,563 reactions, Benjamini-Hochberg (BH) adjusted Wilcoxon p < 0.001 or 0.1, respectively). Many of these reactions are also significantly correlated with the expression of signature genes for Th17 functional activity, which code cytokines and transcription factors [34] (**Figure 2D; Supplementary Figure 2G; Supplementary Table 3**), but not metabolic enzymes. Notably, many classically defined metabolic pathways partially overlapped both with reactions predicted to be pro-pathogenic and reactions predicted to be pro-regulatory (**Figure 2E**), highlighting the value in examining single reactions within a global network rather than conducting a pathway-level analysis. A similar result is obtained at the gene expression level — many metabolic pathways included both genes that were upregulated and genes that were downregulated in Th17p compared to Th17n (**Supplementary Figure 2H, Supplementary Table 4**).

Compass highlighted distinctions in central carbon and fatty acid metabolism between the Th17p and Th17n states, which mirror those found between Th17 and Foxp3^+^ T regulatory (Treg) cells. In central carbon metabolism, Compass predicted that glycolytic reactions, ending with the conversion of pyruvate to lactate are generally more active in the pro-inflammatory Th17p than in the Th17n state **(Figure 2C and 3A)**. This parallels previous results showing that Th17 cells upregulate glycolysis even in the presence of oxygen (hence “aerobic glycolysis”), and that interference with this process promotes a Treg fate [55]–[57]. Compass also predicted an increased activity in Th17p through two segments of the TCA cycle, but not the cycle as a whole (**Figures 2C** and **3A**). A similar breakdown of the TCA cycle in relation with pro-inflammatory function has been shown in macrophages where M1 polarization divided the TCA cycle at the same two points: at isocitrate dehydrogenase (IDH) [58], and at succinate dehydrogenase (SDH) [59], which supported their inflammatory functions [60], [61].

In fatty acid metabolism, Compass predicted that cytosolic acetyl-CoA carboxylase (ACC1), the committed step towards fatty acid synthesis, is upregulated in Th17p, whereas the first two steps of long-chain fatty acid oxidation (long chain fatty acyl-CoA synthetase and carnitine O-palmitoyltransferase (CPT)) were predicted to be significantly higher in Th17n. These predictions mirror a known metabolic difference between the Th17 and Treg lineages, where Th17 cells rely more on *de novo* fatty acid synthesis [62], whereas Tregs scavenge them from their environment and catabolize them and produce ATP through beta-oxidation [56]. We note, however, that recent evidence suggests that CPT may be upregulated in Treg over Th17, but is not functionally indispensable for Treg cells to obtain their effector phenotypes [63].

Among ancillary metabolic pathways, Compass highlighted multiple reactions of amino-acid metabolism that are differentially active between Th17p and Th17n cells **(Figure 2C, Supplementary Table 2)**. It was previously shown that amino acids are important for Th17 cell differentiation [64], and Compass adds further granularity to these findings. In particular, it predicted that serine biosynthesis from 3-phosphoglycerate, as well as three downstream serine fates — sphingosines, choline, and S-adenosyl-methionine (SAM) — were higher in Th17p. On the other hand, parts of urea cycle and arginine metabolism are significantly associated with both pro-Th17p and pro-Th17n states, (**Figure 2C**), suggesting that alternative fluxing within this subsystem may be associated with diverging Th17 cell function. We pursue these predictions and study this subsystem in detail in a companion manuscript (Wang et al., biorxiv preprint). In the following sections we validate the other predictions discussed thus far and build on them to find novel metabolic regulators of Th17 functional states.

### Pathogenic Th17 cells maintain higher aerobic glycolysis and TCA activity, whereas Non-pathogenic Th17 cells oxidize fatty acids to produce ATP

We validated the Compass prediction that pathogenic and non-pathogenic Th17 functional states differ in their central carbon metabolism (**Figure 3a**), using Seahorse assays and liquid-chromatography mass spectrometry (LC/MS) based metabolomics.

**Figure 3.**
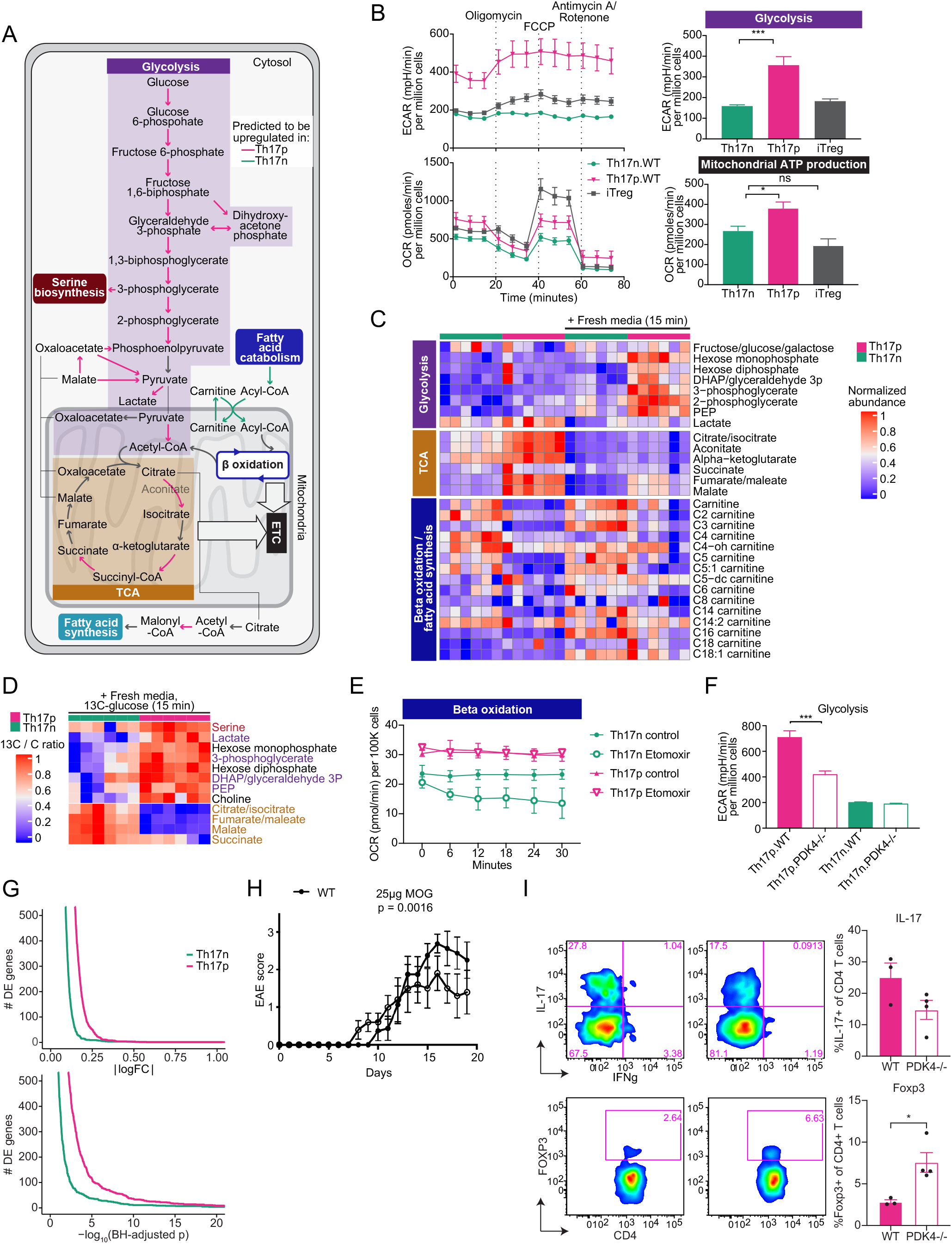
Differential usage of glycolysis and fatty acid oxidation by pathogenic an non-pathogenic Th17 cells. (A) A diagram of central carbon metabolism, overlaid with Compass prediction for differential potential activity between Th17p and Th17n. Differentially active reactions (BH-adjusted Wilcoxon p < 0.1) are colored in magenta (pro-Th17p) or green (pro-Th17n), non-significantly different reactions are colored in grey. (B) Th17n, Th17p and Treg cells were differentiated as described (**Methods**) and replated with Seahorse media at 68h for Seahorse assay. Extracellular acidification rate (ECAR) and oxygen consumption rate (OCR) are reported in response to mitostress test. (C) Th17p and Th17n cells were differentiated and harvested at 68h (left columns) or replated in fresh media with no TCR stimulation or cytokine for 15 minutes (right columns) and subject to LC/MS based metabolomics. (D) Cells were harvested as in C and pulsed with 13C-tagged glucose for 15 minutes. Shown is the ratio of 13C-tagged carbon out of the total carbon content associated with the metabolite (**Methods**). (E) Th17n and Th17p cells were measured for their oxygen consumption rate in the presence of control or 40uM etomoxir. (F) Th17n and Th17p cells from either WT or PDK4-deficient mice were differentiated as described (**Methods**) and replated with Seahorse media at 68h for Seahorse assay. Extracellular acidification rate (ECAR) is reported in response to mitostress test. (G) Number of differentially expressed (DE) genes between PDK4-deficient and WT cells as a function of the significance threshold. (H-I) WT and PDK4-/- mice were immunized with MOG35-55 to induce EAE. (H) EAE clinical score was followed for 21 days. (I) Cells were harvested from CNS at day 15 post immunization for intracellular cytokine or transcription factor analysis.

First, we compared glycolysis and mitochondrial function of Th17p and Th17n cells. A Seahorse assay (which involves culturing cells with glucose-rich media) confirmed that Th17p cells caused significantly higher extracellular acidification (ECAR) than Th17n, indicating accumulation of lactic acid due to aerobic glycolysis **(Figure 3B, top**). Th17p cells also generated significantly more ATP in a mitochondria dependent fashion (**Figure 3B, bottom**), consistent with the predicted higher entrance of pyruvate into the TCA cycle despite the diversion of some pyruvate towards the lactate fate.

Next, we directly measured metabolites within the glycolysis pathway and TCA cycle using LC/MS based metabolomics. When pulsed with fresh media containing glucose (and rested for 15 minutes), there is a substantial increase in glycolytic metabolites in Th17p but less so in Th17n cells (**Figure 3C, top**). Conversely, steady state (pre-pulsing) Th17p and Th17n cells show no apparent difference in glycolytic metabolites, likely due to alternative nutrients or shunting of the glycolytic metabolites into alternative fates. Indeed, after 3 hours with glucose pulsing, the increased level of such metabolites in Th17p return to steady state (**Supplementary Figure 3**).

Interestingly, Compass predicted that two parts of the TCA cycle, but not the cycle as a whole, were upregulated in Th17p: the conversion of citrate to isocitrate and of alpha-ketoglutarate to succinate (mirroring previous findings in macrophages, see above and [58], [59]). LC/MS Metabolomics analysis of cells at steady state revealed that TCA metabolites were generally more abundant in Th17p than in Th17n, apart from succinate **(Figure 3C, middle)**. Therefore, both Compass and the metabolomics data point to succinate as a potential metabolic control point.

To test whether not only absolute metabolite levels, but also the relative allocation of carbon into its possible fates differ between Th17p and Th17n cells, we performed a carbon tracing assay with 13C-glucose. We augmented fresh media with 13C-labeled glucose and computed the ratio of the 13C isotope out of the total carbon for each metabolite. Consistent with our predictions, Th17p had significantly higher relative abundance of 13C-labeled glycolytic metabolites than Th17n (**Figure 3D**). Furthermore, Th17p preferentially incorporated glucose-derived carbon into serine (which branches from glycolysis; **Supplementary Figure 3B**) and its downstream product choline (**Figure 3D**), consistent with shunting of glycolytic metabolites into alternative fates by Th17p cells. This also conforms to the Compass prediction of elevated serine synthesis in Th17p (**Figure 2C**). Th17p cells also had significantly lower relative abundance of 13C-labeled TCA metabolites (**Figure 3D**), suggesting that the higher level of TCA intermediates observed in Th17p at steady state (**Figure 3C**) might not be supported from glucose, but rather from other sources, such as catabolism of amino acids. Taken together, our results suggest that Th17p cells have a higher overall activity through the TCA cycle at steady-state, but quickly switch to aerobic glycolysis when glucose is readily forged from the environment, as observed in the Seahorse, the fresh-media pulsing metabolome assay, and the carbon tracing assay.

We next validated that Th17n cells prefer beta oxidation as predicted by Compass. Metabolomics analysis shows that Th17n cells were enriched in acyl-carnitine metabolites **(Figure 3C, bottom)**, indicative of active lipid transport through the mitochondrial membrane. This could be a result of either increased lipid biosynthesis or increased catabolism (via beta-oxidation), since acylcarnitines are intermediates of both processes. However, acyl-carnitines are noticeably more abundant in Th17n, and short-to medium-length acyl groups are particularly more abundant in the steady state and three hours post glucose pulsing **(Supplementary Figure 3A)**. This supports the hypothesis that under glucose-poor conditions, Th17n cells, more than Th17p, break fatty acids to produce energy (a process which involves the progressive degradation of long-chain fatty acids into shorter acyl-CoA chains, two carbon atoms at a time). Indeed, when etomoxir was used to block acyl-carnitine transportation across mitochondrial membranes, oxygen consumption rate decreased in Th17n but not Th17p cells, as measured by Seahorse assay (**Figure 3E**). Although etomoxir has off-target effects [63], [65], overall our data supports the hypothesis that Th17n cells ultimately divert fatty acid breakdown products into the electron transport chain to generate ATP, which utilizes oxygen as an electron acceptor.

### PDK4-deficient Th17p cells adopt a non-pathogenic-like central carbon program, but retain a pathogenic-like amino acid phenotype

Pyruvate dehydrogenase (PDH) is a critical metabolic juncture through which glycolysis-derived pyruvate enters the TCA cycle (**Supplementary Figure 3B**). Previous studies have shown that the PDH inhibiting kinase 1 (PDK1) is expressed at higher levels in Th17 cells compared to Th1 or FoxP3^+^ Tregs. Consistently, inhibition of PDK1 suppressed Th17 cells but increased the abundance of Tregs [57], whereas PDH activation by PDH phosphatase catalytic subunit 2 (PDP2) had the opposite effects [66]. Compass’s prediction (**Figure 2D**) of increased glycolytic activity, along with these previous studies, prompted us to ask whether PDH has a parallel role in regulating Th17p vs. Th17n states mirroring the reports for Th17 vs. Treg cells [57]. Among PDH inhibitors, PDK4 is of particular interest in immunometabolism, because it plays a role in the cellular starvation response [67], [68].

To determine whether increased glycolysis, regulated by PDH enzymes, in Th17p cells is important for their global metabolic phenotype, we used PDK4^−/−^ mice for perturbation. Despite the low expression of PDK4 mRNA in Th17 cells **(Supplementary Figure 3C)**, Th17 cells differentiated from naive T cells from PDK4^−/−^ mice had reduced conversion of pyruvate to lactate as measured by ECAR in Th17p but not Th17n conditions **(Figure 3F)**. This suggests that PDK4-deficiency increases the alternative pyruvate fate, namely entrance into the TCA only in Th17p cells. It also suggests that Th17n cells control pyruvate entrance to the TCA cycle by other means than PDK4.

To further determine the global transcriptional and metabolic changes induced by PDK4 perturbation, we profiled 146 WT and 132 PDK4^−/−^ Th17p cells and 236 WT and 307 PDK4^−/−^ Th17n cells by scRNA-seq using SMART-Seq2. Consistent with the size of the effects on lactate secretion (**Figure 3F**), we observed a considerably larger effect of PDK4-deficiency on the transcriptome of Th17p cells **(Figure 3G, Supplementary Table 5)**. The genes differentially expressed in Th17p cells were primarily enriched in central-carbon metabolism (**Supplementary Figure 3D** and **Supplementary Table 6**). LC/MS metabolomics showed that PDK4-deficient Th17p cells had notably higher levels of acyl-carnitine, indicating elevated fatty acid transport across mitochondrial membranes (**Supplementary Figure 3E**), similar to WT Th17n cells. Nevertheless, the metabolic phenotype of PDK4-deficient Th17p did not fully shift towards that of Th17n cells. PDK4-deficient Th17p cells retained the WT Th17p metabolome in pathways other than central carbon metabolism, for instance in amino-acid pathways (**Supplementary Figure 3F**). Furthermore, we did not observe significant differences in the expression of key cytokines or transcription factors that are associated with the effector function of these cells.

Seeing that PDK4-deficiency had partially shifted Th17p central carbon metabolism towards the Th17n state *in vitro*, we next tested the pertaining effects *in vivo*. To this end, we studied the impact of PDK4-deficiency on the development of EAE, an autoimmune disease induced by pathogenic Th17 cells. Consistent with previous studies that glycolysis promotes inflammation [57], [69]–[71], mice with global knockout of PDK4 developed less severe disease as determined by the clinical disease scores **(Figure 3H)** with decrease in Th17 cells and increase in the infiltration of Foxp3^+^ Tregs in the CNS of the mice undergoing EAE (**Figure 3I**). We therefore conclude that PDK4-deficient Th17p cells resemble Th17n in their central carbon metabolic state, but not in other metabolic pathways. These results prompted us to interrogate metabolic differences outside of central carbon pathways, which are presented in the companion manuscript (Wang et al., biorxiv preprint).

### The glycolytic enzyme phosphoglycerate mutase (PGAM) suppresses Th17 cell pathogenicity

Thus far, our analysis relied on an *inter-population* comparison between the extreme states of Th17n and Th17p cells. However, we have previously shown that there is also considerable continuous variation in the transcriptomes of Th17n cells, which spans into pathogenic-like states [29]. To explore the relationship between metabolic heterogeneity and pathogenic potential within the Th17n subset, we next performed an intra-population analysis of Th17n cells. This also demonstrates that Compass can be applied to scRNA-Seq data in cases where the states of interest (e.g., Th17n vs. Th17p) are either unknown or cannot be experimentally partitioned into discrete types. To perform an intra-population Compass analysis of single Th17n cells, we correlated the Compass scores associated with each reaction with the pathogenicity gene signature scores of the respective cells **(Methods)**.

While the resulting correlations of individual reactions with the pathogenicity score were largely consistent with the results of the inter-population analysis (Th17p vs. Th17n) **(Figure 4A, Supplementary Tables 7-8**), the intra-population analysis predicted that some glycolytic reactions may be negatively, rather than positively (as in the inter-population analysis), associated with Th17 pathogenicity. The most notable of these reactions was the one catalyzed by the enzyme phosphoglycerate mutase (PGAM), which was negatively associated with pathogenicity in the intra-population analysis of Th17n cells **(Figure 4A)**, but positively associated with Th17p cells in the inter-population analysis **(Figure 2B,C)**. This prediction was unexpected because increased glycolysis is generally known to support pro-inflammatory phenotypes in T cells (see also **Discussion**) [55]–[57], [69], [72]–[78].

**Figure 4.**
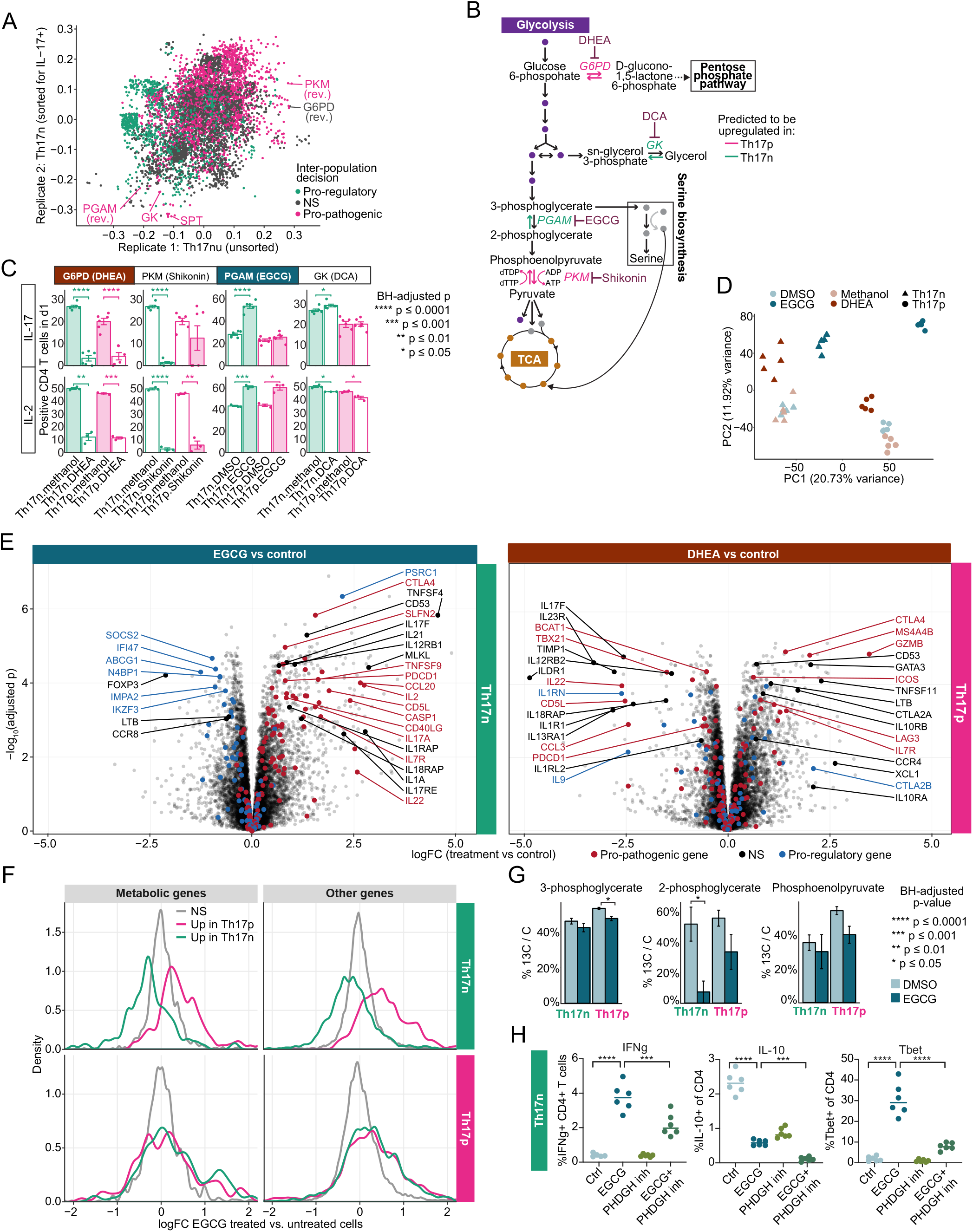
An unexpected role for PGAM in mediating TGFb-induced Th17 pathogenicity. (A) Intra-population analysis in two biological replicates (the Th17n and Th17nu cell populations, see **Figure 2a**). Dots are single metabolic reactions, and axes denote their correlation with the pathogenic signature in the Th17nu and Th17n groups. Colors denote whether the reaction was decided as pro-inflammatory, pro-regulatory, or non-significantly (NS) associated with either state by the inter-population analysis. PGAM, GK, PKM, and G6PD are reactions discussed in the manuscript (see Figure 4b). SPT = serine-pyruvate transaminase (EC 2.6.1.51). Rev = reverse (backwards) direction. (**Methods**). (B) Schematics of central carbon metabolism, the highlighted magenta and green reactions are the two predicted to be most correlated and anti-correlated with the computational pathogenicity score within the Th17n compartment, respectively. Reported inhibitors of these reactions are denoted. (C) Effects of inhibiting candidate genes on Th17 cytokines as measured by flow cytometry are shown. Naïve T cells were differentiated under pathogenic (Th17p) and non-pathogenic (Th17n) Th17 cell conditions (**Methods**) in the presence of control solvent or inhibitors. Cells were pre-labeled with division dye and protein expression is reported for cells that have gone through one division (d1) to exclude arrested cells. (D) PCA of bulk RNA-Seq of d1 Th17 cells as in B. (E) Differential gene expression due to EGCG and DHEA treatment. Red and blue dots represent genes associated with the pro-pathogenic and pro-regulatory Th17 transcriptional programs, respectively (red genes are ones belonging either to the list of pro-pathogenic Th17 markers (Figure 2d and **Methods**) or to the Th17 pro-inflammatory covariation module defined by [29]; blue genes are similarly defined). Highly differential genes associated with surface receptors, cytokine activity, or that are otherwise of interest are labelled by name. (F) Histograms of the logFC per gene in differential expression of EGCG-vs. DMSO-treated cells. A separate histogram is shown for Th17p-associated (magenta), Th17n-associated (green), and non-significantly associated (grey) genes. Genes were partitioned into these three groups by differential expression in bulk RNA-Seq (same libraries as shown in panel D) between DMSO-treated Th17p and Th17n cells with significance threshold of BH-adjusted p ≤ 0.05 and log2 fold-change ≥ 1.5 in absolute value. (G) ratio of 13C-tagged carbon to total carbon in Th17 cells cultured for 15 minutes in the presence of 13C-glucose. Three metabolites are shown: PGAM’s substrate (3-phosphoglycerate), product (2-phosphoglycerate), and the next downstream metabolite along the glycolytic pathway (phosphoenolpyruvate). (H) Th17n cells were differentiated in the presence of solvent alone, EGCG, PHDGH inhibitor (PKUMDL-WQ-2101, **Methods**), or the combination. Cells were harvested at 96h for flow cytometric analysis.

To functionally validate the glycolytic targets associated with Th17 cell pathogenicity by the intrapopulation analysis, we used chemical inhibitors against enzymes driving the top two glycolytic reactions that were most positively correlated (regulated by pyruvate kinase muscle isozyme [PKM], and glucose-6-phosphate dehydrogenase [G6PD]) and top two that were most negatively correlated (phosphoglycerate mutase [PGAM], and glucokinase [GK]) with the pathogenicity score (**Figure 4B**). The inhibitors were shikonin (inhibits PKM2), dehydroepiandrosterone (DHEA, inhibits G6PD), epigallocatechin-3-gallate (EGCG, inhibits PGAM1), and 2,3-dihydroxypropyl-dichloroacetate (DCA, inhibits GK) (**Methods**).

We first analyzed the effects of inhibitors on Th17n and Th17p cell differentiation and function using flow cytometry (**Figure 4C**). Due to the possibly deleterious effects of blocking these central reactions on cell viability, we used the highest dose of each inhibitor that did not affect cell viability (compared to solvent alone). We further used flow cytometry to restrict the analysis to cells that had undergone one division (d1) so as to exclude arrested cells or cells that have been blocked from activation and expansion. In addition, since two different solvents (DMSO and methanol) were needed for different inhibitors, every treatment group was matched with an appropriate vehicle control. We found that IL-17 expression conformed to the prediction made by Compass. It was significantly upregulated by chemical inhibition of the two enzymes (PGAM or GK) predicted to suppress pathogenicity, and downregulated by chemical inhibition of the two enzymes (G6PD or PKM) predicted to promote pathogenicity **(Figure 4C)**. This was further confirmed when profiling a larger set of cytokines secreted by Th17 cells: inhibition of PKM or G6PD curtailed all cytokine production suggesting that these enzymes are important for overall T effector functions. In contrast, cells with PGAM or GK inhibition, at the optimal concentration, mostly retained their cytokine profile with a few exceptions **(Supplementary Figure 4)**.

To analyze the impact of perturbing glycolytic enzymes on the transcriptome, we used bulk RNA-Seq to profile Th17n and Th17p cells grown in the presence of either the predicted pro-regulatory inhibitor DHEA (inhibiting G6PD) or the predicted pro-inflammatory inhibitor EGCG (inhibiting PGAM) (**Figure 4D-F**). The first principal component (PC1), which is the main axis of variation in the data, represented as expected, the pathogenicity phenotype. In both Th17n and Th17p cells, EGCG shifted cell profiles towards a more pathogenic state on PC1, whereas DHEA shifted them to a less pathogenic state (**Figure 4D**). The difference between the two vehicle controls was inconsequential compared to cell type and interventions.

To better interpret the drug-induced transcriptional changes, we examined individual genes whose expression is associated with either Th17n or Th17p effector function as wells as global transcriptomic shifts **(Figure 4E,F** and **Supplementary Table 9)**. A comparison of DHEA to vehicle control identified a large number of effector genes that are modulated **(Figure 4E, right)**. These include a significant decrease in *IL23R* and *TBX21* transcripts in both Th17p and Th17n, two genes critical for Th17 cell pathogenicity, and in IL9 and IL1RN, two genes highly expressed in non-pathogenic Th17 cells (Lee et al. 2012). Conversely, EGCG clearly strengthened the pathogenic transcriptional program in Th17n, globally upregulating pro-inflammatory genes (e.g., IL22, IL7R, and CASP1) and (to a more limited extent) downregulating pro-regulatory ones (e.g., IKZF3) **(Figure 4E, left and 4F)**. The global shift towards the pro-inflammatory Th17 program was observed both in metabolic and non-metabolic transcripts, supporting the hypothesis that PGAM inhibition by EGCG effected a network-wide metabolic shift that mediated emergence of a pro-inflammatory Th17 program **(Figure 4F)**.

To verify that the effect of EGCG was mediated by a specific inhibition of PGAM (rather than an off-target effect) we conducted a carbon tracing assay in which the cell’s medium was supplemented with 13C-glucose **(Methods)**. PGAM inhibition with EGCG led to a sharp decrease (from 51% 13C ratio to 7% in Th17n and from 55% to 33% in Th17p) in 13C contents of 2PG (PGAM’s product) but not 3PG (PGAM’s substrate) or any other glycolytic metabolite that we were able to measure **(Figure 4G)**. Interestingly, 13C ratio of PEP (one step downstream of 2PG) was not changed as well. This suggests that the effect of the inhibitor is restricted (at least within glycolysis) to the PGAM reaction that lies directly downstream of 3PG.

As the serine biosynthesis pathway is more active in Th17p than in Th17n (**Figure 2C**) and lies directly downstream of 3PG (**Figure 4B**), we asked whether inhibiting serine biosynthesis can rescue the effect of PGAM inhibition. To this end, we treated Th17n cells with inhibitors to PGAM and PHGDH (phosphoglycerate dehydrogenase), alone or in combination. We found that further inhibiting PHDGH rescued the upregulation of Tbet and IFNg induced by EGCG but not its impact on IL-10 suppression (**Figure 4H**).

Taken together, an intra-population Compass analysis predicted that within the Th17n compartment, the glycolytic PGAM reaction inhibits, rather than promotes, pathogenicity. This prediction relied on heterogeneity within the Th17n population, yielding results that are contrary to those from inter-population comparisons of Th17 to Treg or Th17p to Th17n. EGCG specifically inhibited this reaction, and promoted a transcriptional state indicative of a more pro-inflammatory potential, as evidenced by a global shift in the transcriptome toward a Th17p-like profile. RNA-Seq further supported the hypothesis that EGCG mediates its effects by altering the cellular metabolic profile.

### PGAM inhibition exacerbates, whereas G6PD inhibition ameliorates, Th17-mediated neuroinflammation *in vivo*

To test the functional relevance of the transcriptome shifts induced by EGCG and DHEA *in vivo*, we used the adoptive T cell transfer system, so that the effect of inhibitors is limited to T cells rather than all cells in the host. We generated Th17n and Th17p cells from naive CD4+ T cells isolated from 2D2 TCR-transgenic mice, with specificity for MOG 35-55, and transferred them into wildtype mice to induce EAE.

Consistent with Compass prediction, Th17p cells treated with DHEA reduced the severity of disease at peak of EAE in the recipients (**Figure 5A**). By the time the mice were sacrificed, however, the number of lesions in CNS was not significantly different (**Figure 5B**), and surviving mice showed no significant alterations in antigen-specific cytokine secretion in response to MOG, except for increased IL-2 in the DHEA treated group (**Supplementary Figure 5A**). More interestingly, and in agreement with Compass predictions, EGCG-treated Th17n cells induced EAE, albeit in mild form, whereas solvent treated cells failed to produce any consequential neuroinflammation **(Figure 5C)**. Recipients of EGCG-treated Th17n cells had a significantly higher EAE incidence rate (10/12) compared with the control group (0/12, Fisher’s exact p = 1.1*10^−4^). Consistent with the clinical disease, histological analyses revealed an increased number of CNS lesions in in both the meninges and the parenchyma of mice that were injected with EGCG-treated Th17 cells (**Figure 5D**). While there was only a small difference in antigen-specific T cell proliferation (**Figure 5E**), there was a significant increase in secretion of IL-17, IL-17F, IL-22 and IL-6 (**Figure 5F** and **Supplementary Figure 5B**) in response to antigen in cells isolated from draining lymph node of mice transferred with EGCG-treated Th17n cells. As EGCG treated non-pathogenic Th17n cells induced only mild EAE, we asked whether EGCG will further enhance encephalitogenicity of Th17 cells if IL-23 is included in the differentiation cultures, which stabilizes the Th17 phenotype [79]–[82]. IL-23-treatment indeed enhanced EAE disease severity, but still Th17n cells treated with IL-23+EGCG induced significantly more severe EAE than their IL-23+solvent-treated counterparts (**Figure 5G**). Histopathology across all experiments revealed that EGCG treatment of Th17n cells promoted, whereas DHEA treatment Th17p cells restricted, optic neuritis/perineuritis in host mice (**Figure 5H**). Interestingly, mice transferred with EGCG-treated Th17 cells (Th17n or Th17n with IL-23) were the only experimental group to produce Wallerian degeneration in proximal spinal nerve roots (**Figure 5I, J)**.

**Figure 5.**
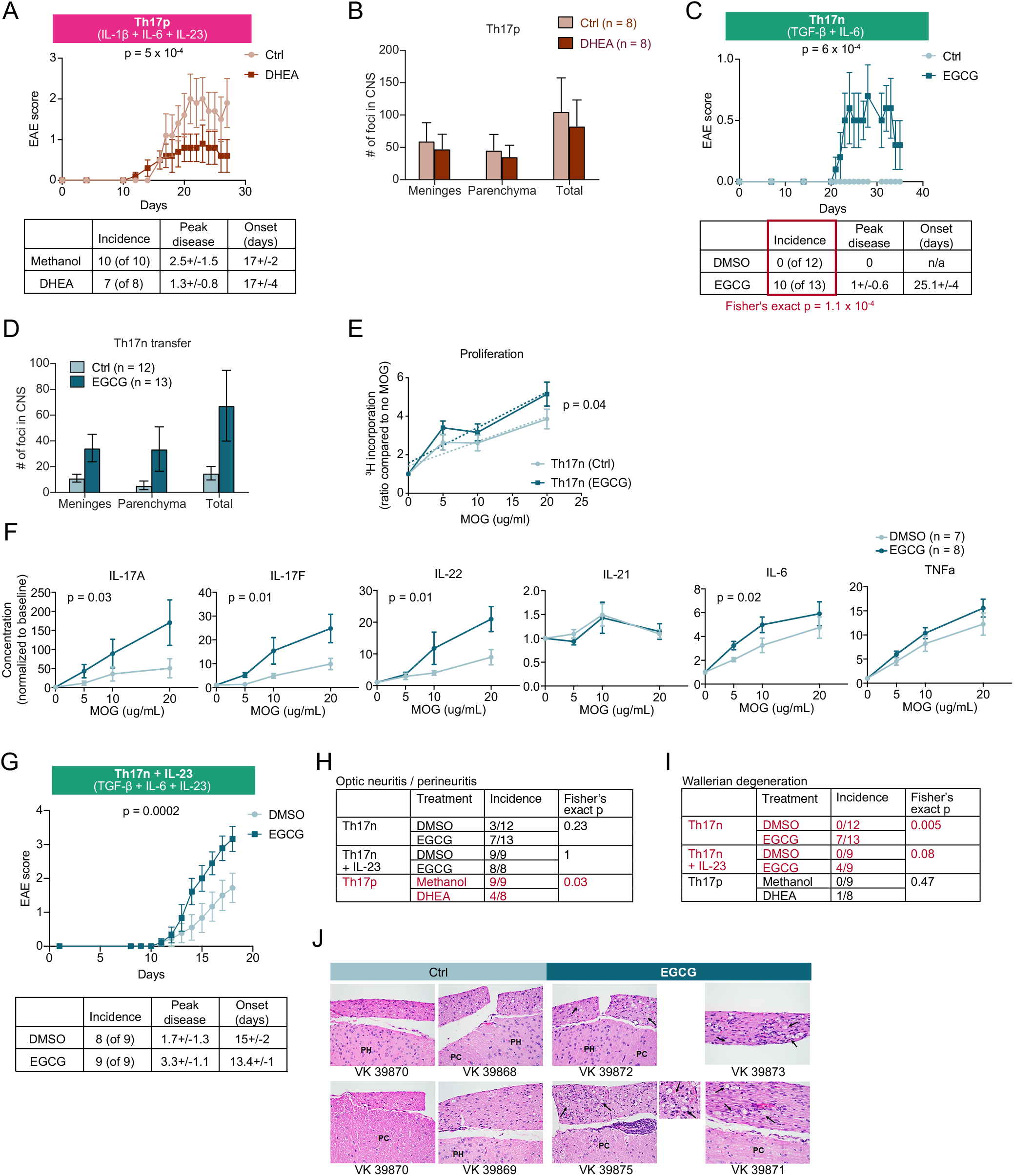
EGCG exacerbates and DHEA ameliorates Th17-induced EAE *in vivo*. 2D2 TCR–transgenic Th17 cells were adoptively transferred after differentiation in vitro in the presence of an inhibitor or vehicle as indicated. (A, C, G) Clinical outcome of EAE; (B, D) Histological score based on cell infiltrates in meninges and parenchyma of CNS; (E, F) Draining lymph node (cervical) from respective mice were isolated and pulsed with increasing dose of MOG_35-55_ peptide for 3 days and (E) subjected to thymidine incorporation assay; or (F) measurement of cytokine secretion by Legendplex and flow cytometry. Concentrations were normalized through division by the respective response to no antigen control (H-I) Independent pathological report of CNS isolated from mice with EAE at end point (d35 for EGCG experiments; d28 for DHEA experiment); Optic nerves were not found in the histologic section from one animal in the EGCG+IL-23 group. (J) Representative histology of spinal cord and spinal nerve roots. There is greater meningeal inflammation and Wallerian degeneration (digestion chambers, arrows) in posterior spinal nerve roots in EGCG vs. Control mice. PC, posterior column; PH, posterior horn. Individual mouse numbers are indicated. The smaller panel shows VK 39875 mouse section at higher magnification. All are H. & E., 40X objective. Three similar experiments were performed.

In conclusion, Compass correctly predicted metabolic targets including glycolytic pathways whose deletion affected Th17 function. Importantly, it was able to pinpoint a glycolytic reaction that suppresses Th17 pathogenicity, which runs contrary to the current understanding that aerobic glycolysis as a whole is associated with a pro-inflammatory phenotype in Th17 cells.

## Discussion

We presented Compass — a flux balance analysis (FBA) algorithm for the study of metabolic heterogeneity among cells based on single-cell transcriptome profiles and validated a number of predictions by metabolome and functional analyses. Compass successfully predicted metabolic targets in both central and ancillary pathways based on its network approach. These results support the power of transcriptomic-based FBA to make valid predictions in a mammalian system.

Glycolysis is a central regulator of T cell function. Compass predicted an association between aerobic glycolysis and Th17 pathogenicity, which accords with multiple previous results tying elevated glycolysis with T cell inflammatory functions. However, a Compass-based data-driven analysis based on scRNAseq unexpectedly revealed that not all glycolytic reactions promote the pro-inflammatory phenotype in Th17 cells. This result was obtained via an intra-population analysis of individual cells. It serves as a further example to the power of studying single-cell heterogeneity within seemingly homogenous populations (here, Th17n), which allowed us to identify a novel regulator that would have otherwise been missed at a population level (here, a comparison of Th17p and Th17n). The computational prediction and the data corroborating it also demonstrate that despite the common assumption that glycolysis promotes pro-inflammatory functions in Th17 cells and other immune compartments [2], [3], [83]–[87], the role of glycolysis in induction of pro-inflammatory phenotypes may more nuanced [88], [89].

Static FBA algorithms assume that the system under consideration operates in chemical steady state [90]. Even under this assumption, there remains an infinite number of feasible flux distributions that satisfy the preset biochemical constraints. Therefore, most studies assume that the system (here, a cell) aims to optimize some metabolic function, usually production of biomass or ATP [91]. However, whereas such objectives may successfully predict phenotypes of a unicellular organism [92], they are ill-suited for studying mammalian cells [93]. To overcome this challenge, rather than optimizing a single metabolic objective function, Compass optimizes a set of objective functions, each estimating the degree to which a cell’s transcriptome supports carrying the maximal theoretical flux through a given reaction. The result is a high dimensional representation of the cell’s metabolic potential (one number per reaction). A biological signal (e.g., differences in reaction potential) can be detected in this high-dimension owing to the statistical power afforded by the large number of cells in a typical scRNA-Seq dataset. Nonetheless, there is no inherent limitation preventing one from applying Compass to study bulk (i.e., non-single-cell) transcriptomic data.

The database of metabolic reactions we used pertains to human cells, and as such our study does not address differences between human and mouse metabolism. In addition, the database provides a global view of the metabolic capabilities of a human cell, accrued from various sources and in diverse cell types. Not all reactions may be functional in a studied cell type, or under particular physiological conditions. This concern can be addressed to some extent by procedures for deriving organ-specific metabolic models [94]. Moreover, the metabolic state of a cell depends on the nutrients available in its environment, which are often poorly characterized. Here, our computations assume an environment rich with nutrients, which accords with the studied *in vitro* growth media. Modifying this to better represent physiological conditions should increase the algorithm’s predictive capabilities, especially for cells derived *in vivo*, where nutrient scarcity may be a limiting factor, and nutrient availability may vary between tissues.

One of the outstanding challenges in the field of single cell genomics is translating the vast data sets presented in cell atlases into an actionable knowledge resource, i.e. using observed cell states to deduce molecular mechanisms and targets [16]. Compass was designed with this challenge in mind, and addresses it in the metabolic cellular subsystem, which can be tractably modeled *in silico*. In light of the wide appreciation of cellular metabolism as a critical regulator of physiological processes in health and disease, we expect Compass to be useful in predicting cell metabolic states, as well as actionable metabolic targets, in diverse physiological and pathologic contexts.

## Code and data availability

We are working on making an open-source and free for academic use implementation available on Github, and will update this preprint when it is available. In the meantime, code is available upon request. A GEO deposition of RNA-Seq data is underway as well.

## Author Contributions

AW, CW, AR, VKK, and NY conceived and designed the study. AW, DD, and NY designed the algorithm and performed computational analyses. DD and SE developed software packages. CW designed, performed, and analyzed experiments, with significant contributions by JAP, KP, and CC in LC/MS metabolomics, NRH and MH in Seahorse assays, and further assistance by SZ, JF, EAG and VPD. VKK and NY supervised the study. AW, CW, VKK, and NY wrote the manuscript with contributions from all authors. All authors read and approved the final manuscript

## Acknowledgments

We thank Eytan Ruppin and Michael B. Cole for fruitful discussions and Christina Usher for artwork. NY and AW were supported by the Chan Zuckerberg Biohub and by a National Institute of Mental Health (NIMH) grant NIH5U19MH114821. This work was supported by grants from the National Institutes of Health (RO1NS30843, R01AI144166, R01NS045937, P01AI073748, P01AI039671, P01AI056299) awarded to VKK, and by a Career Transitional Fellowship from the National Multiple Sclerosis Society awarded to CW. JF was supported by a Max Kade fellowship awarded by the Austrian Acadamy of Science (ÖAW).

---

**Supplementary Figure 1.**
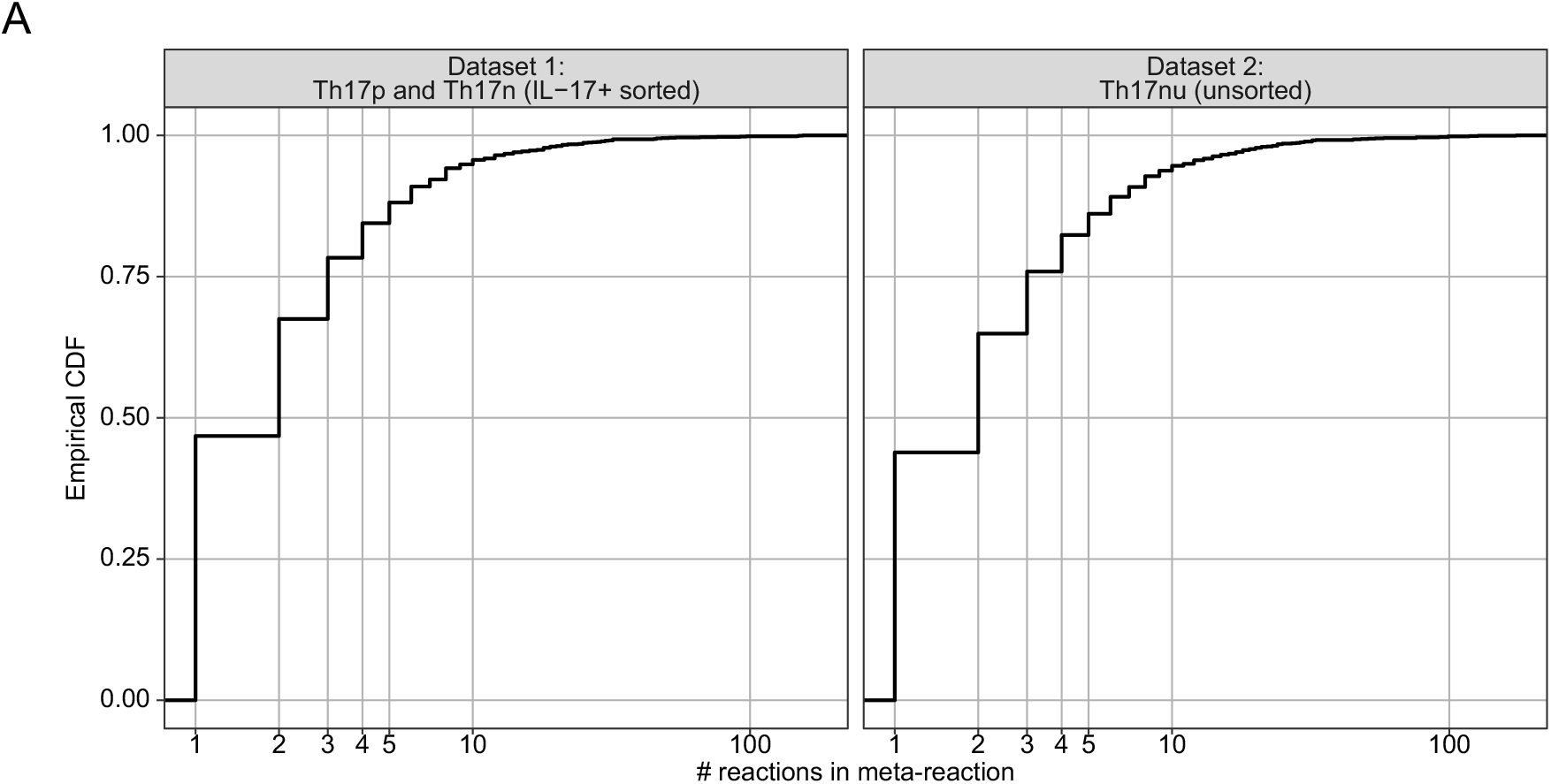
Cumulative distribution function (CDF) of number of reactions per meta-reaction.

**Supplementary Figure 2.**
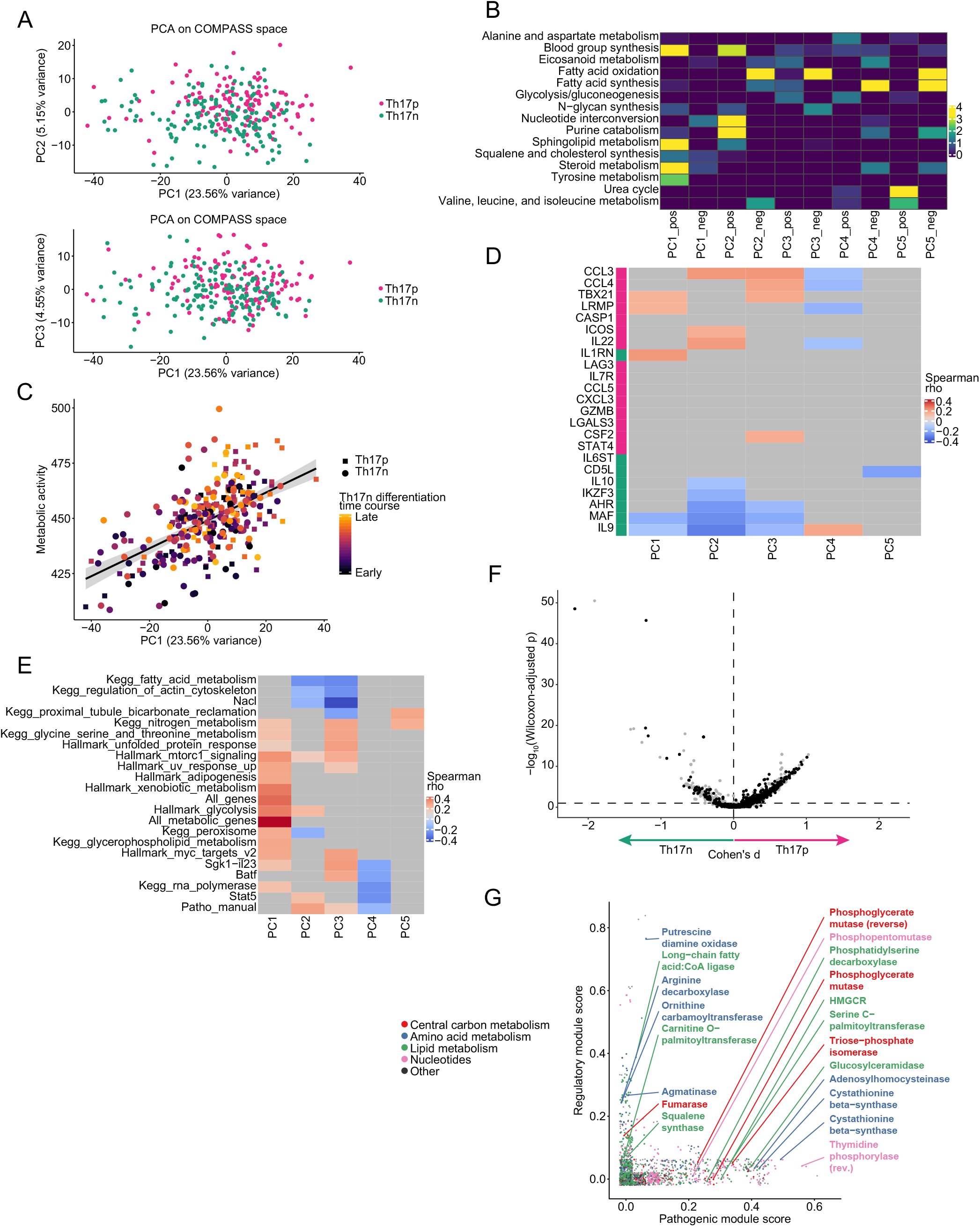
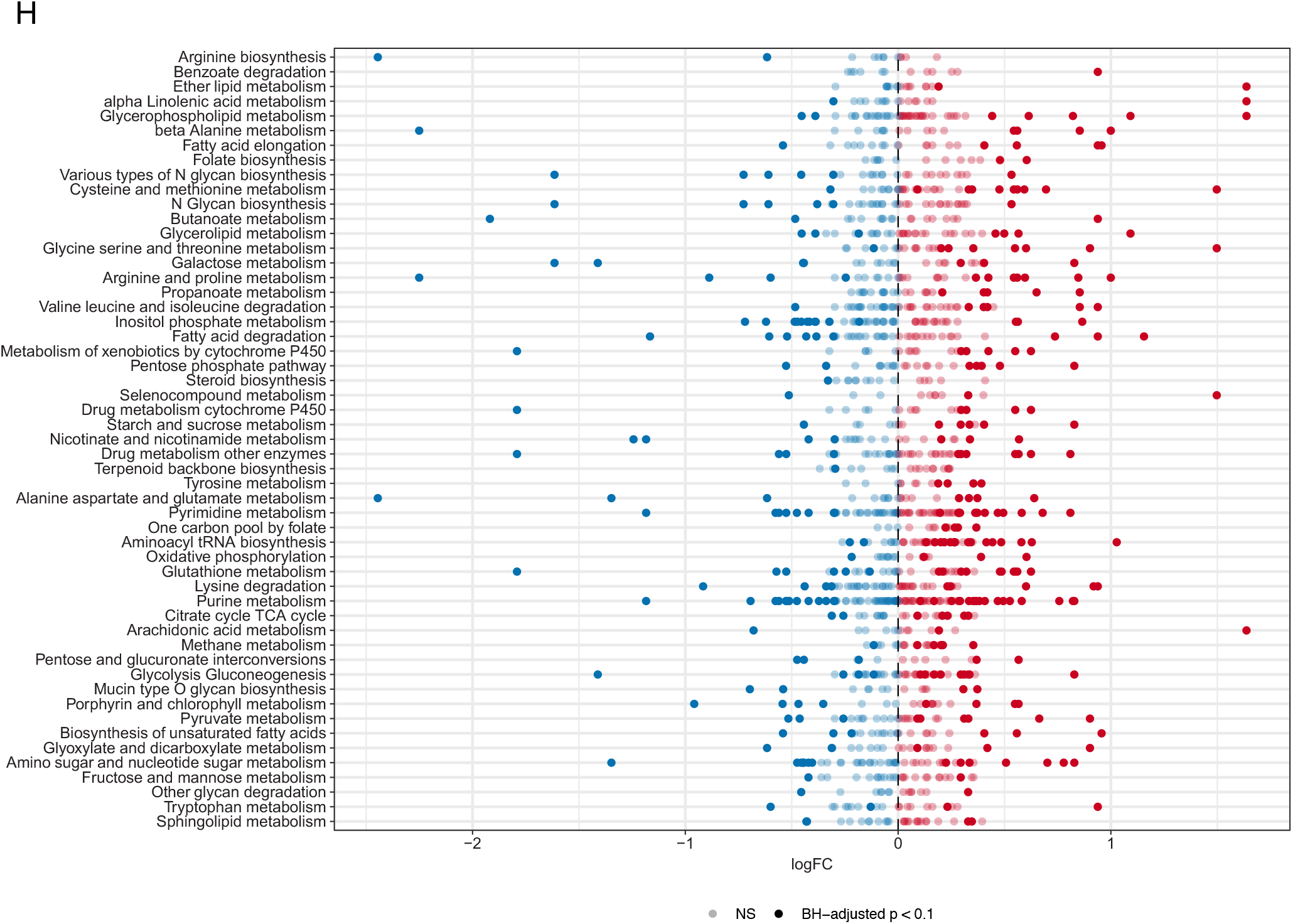
(A-E) PCA of Compass space restricted to core meta-reactions, see main text. (A) PC1 scores plotted against PC2 and PC3 scores. (B) Enrichment of metabolic pathways in the positive or negative directions of top principal components. Enrichment is computed with GSEA [95] over single reactions (rather than genes, as in the common applications). Colors are −log10(BH-adjusted p), truncated at 4, with p being the GSEA p value. Pathways correspond to Recon2 subsystems. (C) PC1 scores plotted against computational signatures of cellular metabolic activity and Th17 differentiation time course (**Methods**). (D) Spearman correlation of top PCs with known pro-pathogenic (magenta) and pro-regulatory (green) marker genes, none of which is metabolic. Only significant correlations (BH-adjusted p < 0.1) are shown in color. (E) Spearman correlation of computational transcriptome signatures with the top principal components. Only significant correlations (BH-adjusted p < 0.1) are shown in color and non-significant correlation coefficients are greyed out. See **Methods** for signature computation. (F) Same analysis as shown in Figure 2c, but showing all reactions (and not just ones belonging to certain pathways, as in the main figure). (G) We computed a pro-pathogenic score for each reaction by taking the ratio of pro-pathogenic and pro-regulatory markers with which it correlates and anti-correlates, respectively (BH-adjusted p < 0.1 for a Spearman correlation) out of the 23 marker genes (listed in **Figure 2d** and **Methods**). Similarly, we computed pro-regulatory reaction scores. Only core reactions are shown. (H) Same analysis as shown in Figure 2E, only at the gene expression level (and not reaction level based on Compass scores). Genes are grouped by KEGG pathways (and may be annotated as belonging to more than one pathway).

**Supplementary Figure 3.**
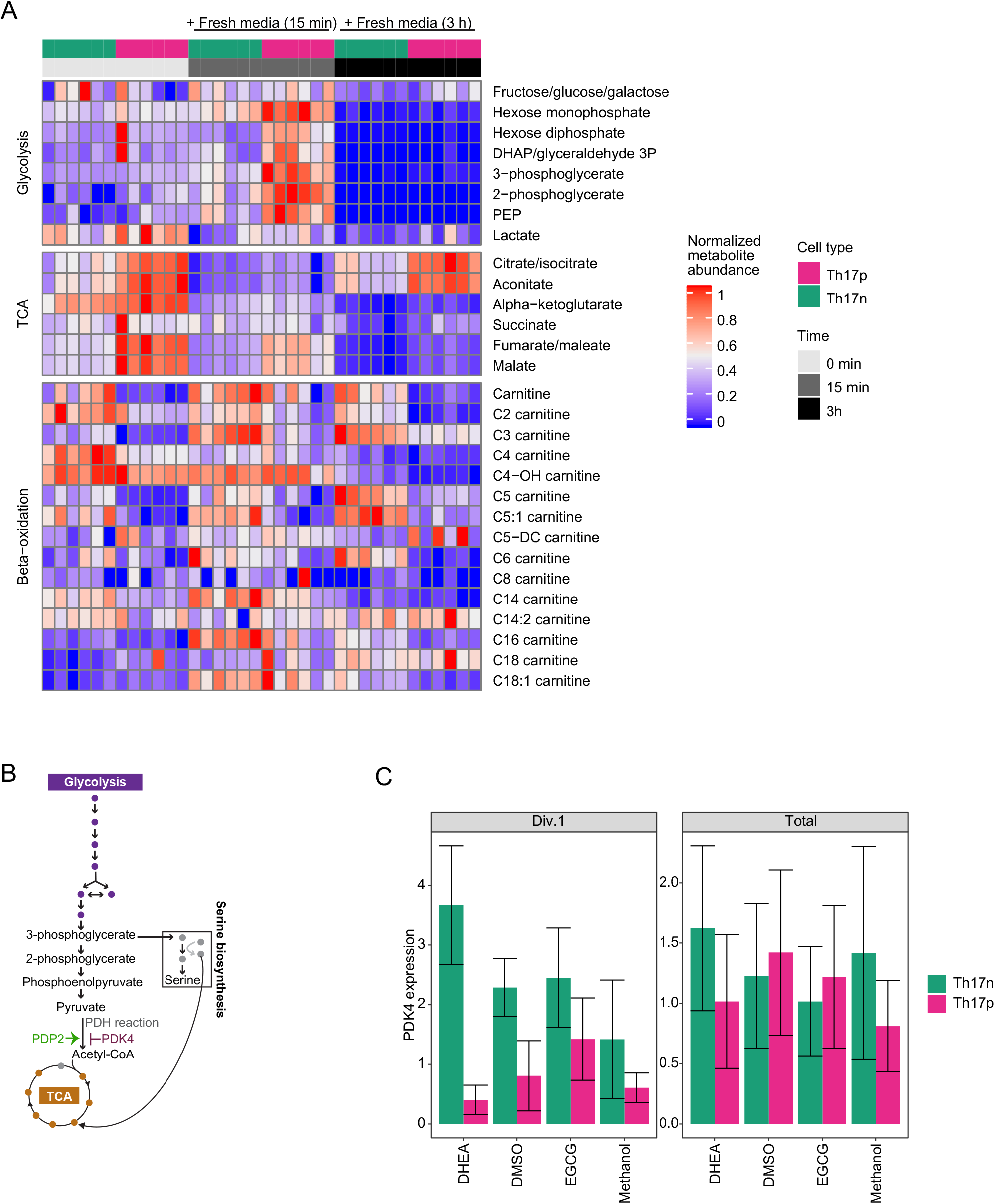
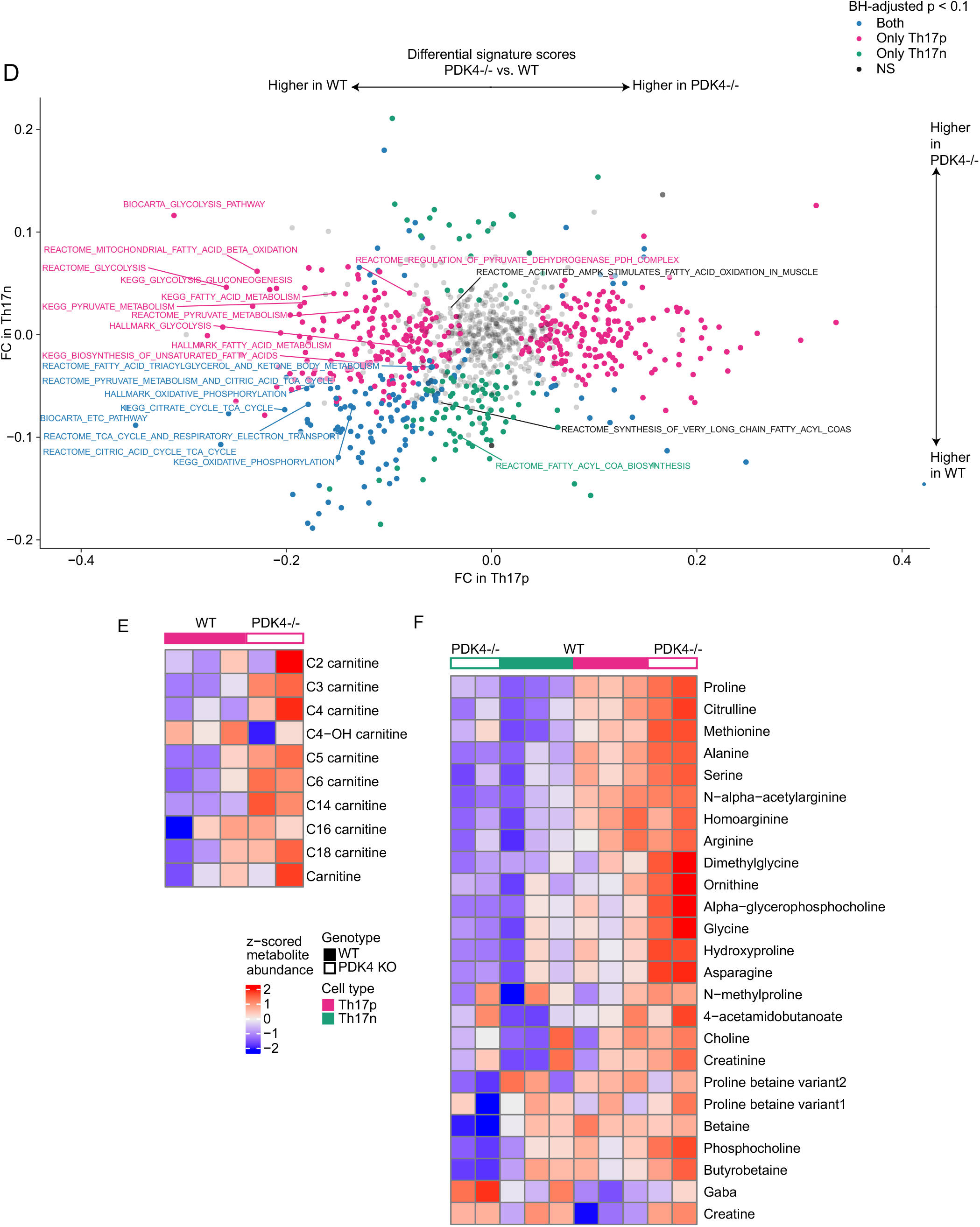
(A) Parallel of main figures 3c showing also 3h after fresh media pulse. (B) The glycolysis pathway, as shown in main figure 4a, highlighting PDH and associated reactions. (C) PDK4 transcript expression in the experiment described in main Figure 4c. (D) Dots are transcriptomic computational signatures (**Methods**), axes correspond to the fold-change in the signature’s value in comparisons of PDK4-deficient cells vs. WT cells in Th17p (x-axis) and Th17n (y-axis). (E) Th17 cells from PDK4-/- and WT mice were subject to LC/MS metabolomics as in main Figure 3c, having been replated for 15 minutes. (F) metabolites associated with amino-acid metabolic pathways in the assay described in main Figure 3c.

**Supplementary Figure 4.**
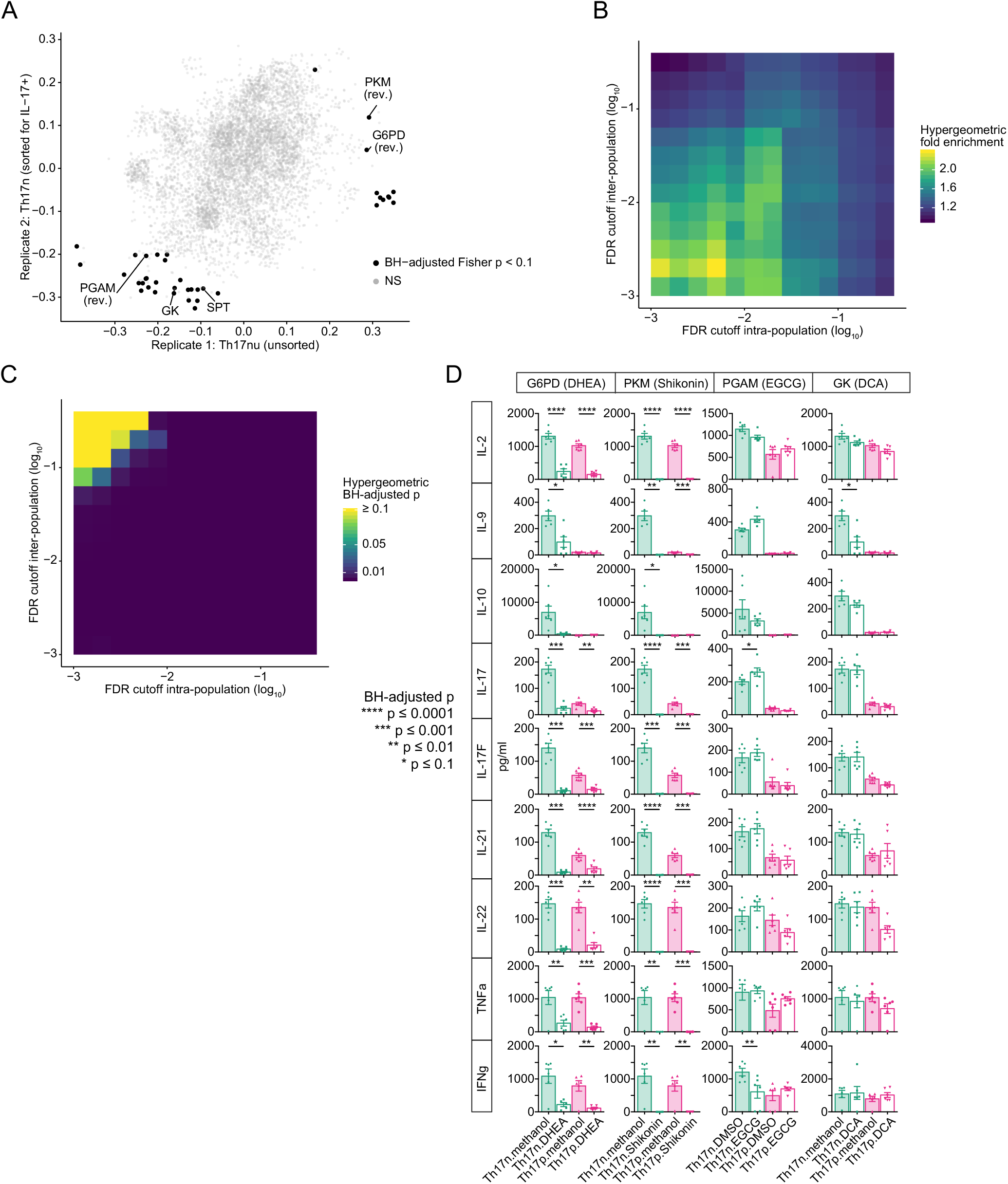
(A) Same data as shown in Figure 4a, highlighting the reactions with significant adjusted Fisher p value in the intra-population analysis; every reaction is assigned a combined Fisher p-value of the two p-values measuring the significance of the correlation with the two axes (**Methods**). Search space was limited to core reactions. (B-C) Hypergeometric enrichment of the targets identified by the inter-population analysis (reactions with differential potential activity between Th17p and Th17n, decided by a BH-adjusted p cutoff) in targets identified by the intra-population analysis (reactions identified by a BH-adjusted Fisher p cutoff) while varying the cutoffs. (D) Supernatant from Th17 cell cultures performed for main Figure 4c are harvested for cytokine analysis using Legendplex.

**Supplementary Figure 5.**
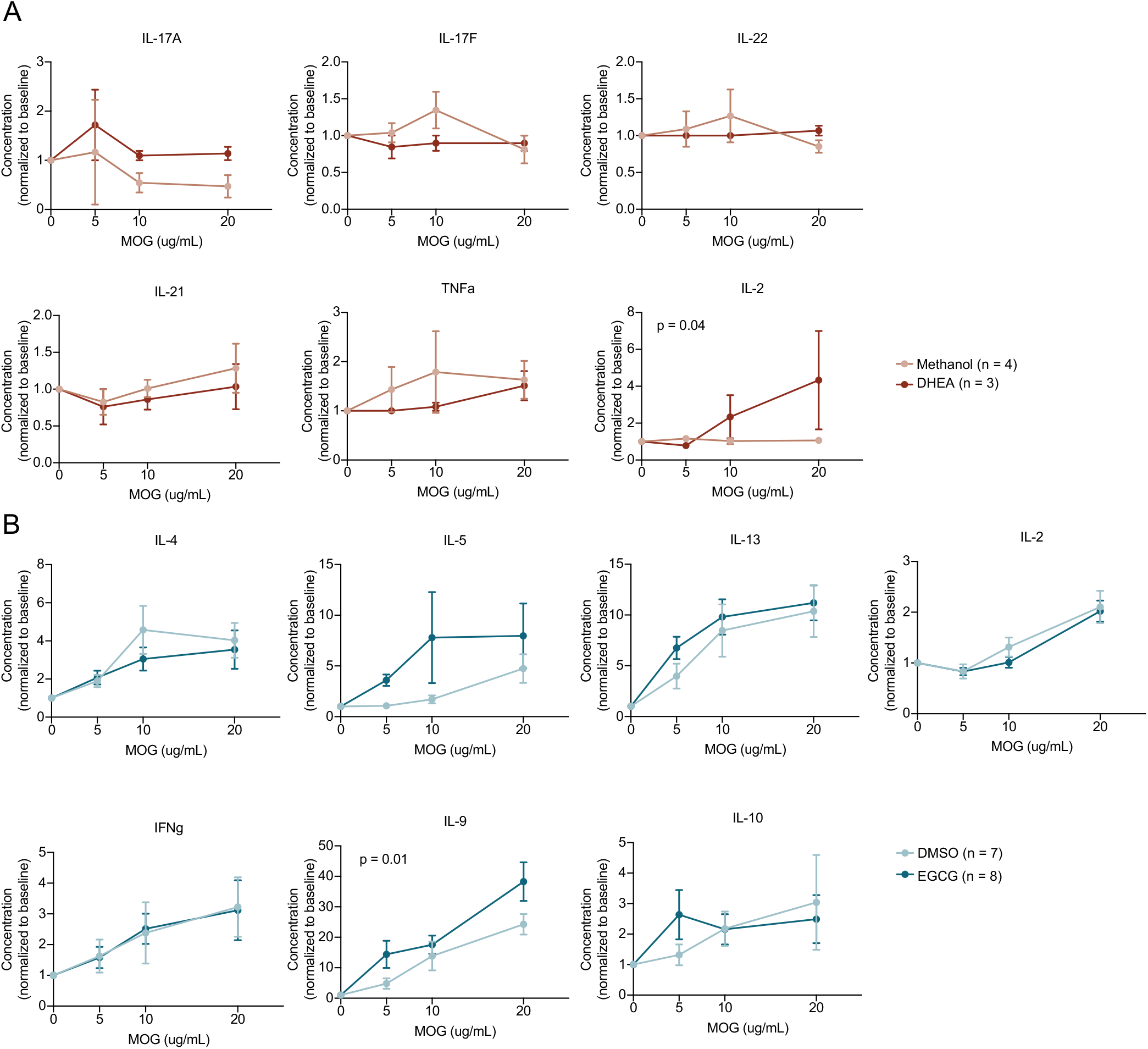
Cytokine secretion after three days of culture with increasing dose of MOG35-55 peptide from cells isolated from draining lymph node (cervical) of mice transferred with (A) methanol or DHEA treated Th17p cells as in Figure 5A or (B) DMSO or EGCG as in Figure 5C. Concentrations were normalized through division by the respective response to no antigen control.

